# Identification, analysis, and confirmation of seed storability-related loci in Dongxiang wild rice (*Oryza rufipogon* Griff.)

**DOI:** 10.1101/2021.11.16.468766

**Authors:** Minmin Zhao, Biaolin Hu, Yuanwei Fan, Gumu Ding, Wanling Yang, Yong Chen, Yanhong Chen, Jiankun Xie, Fantao Zhang

**Affiliations:** College of Life Sciences, Jiangxi Normal University, Nanchang 330022, China; Rice Research Institute, Jiangxi Academy of Agricultural Sciences, Rice National Engi-neering Laboratory, Nanchang 330022, China; Jiangxi Provincial Key Lab of Protection and Utilization of Subtropical Plant Resources, Nanchang 330022, China; College of Biological Sciences and Biotechnology, Beijing Forestry University, Beijing 100083, China; Department of Biology and Center for Engineering Mechanobiology, Washington University in St.Louis, St.Louis, MO 63130, USA

**Author notes:** Correspondence (F.Z.); (J.X.). Contributed equally to this work (Minmin Zhao, Biaolin Hu, Yuanwei Fan). (M.Z.); (G.D.); (Y.H.C.); (B.H.); (W.Y.); (Y.C.); (Y.F.).

**Keywords:** wild rice, seed storability, BSA-seq, genetic resource, rice improvement

## Abstract

Dongxiang wild rice (*Oryza rufipogon* Griff.) (DXWR) has strong seed storability and identifying its elite gene resources may facilitate genetic improvements in rice seed storability. In this study, we developed two backcross inbred lines (BILs) populations, with DXWR as a common donor parent and two rice varieties (F6 and R974) as recipient parents. Bulked segregant analysis via whole genome sequencing (BSA-seq) was used to identify seed storability-related loci in the DXWR and F6 population. Two main genomic regions containing 18,550,000–20,870,000 bp on chromosome 4 and 7,860,000–9,780,000 bp on chromosome 9 were identified as candidate loci of DXWR seed storability; these overlapped partially with seed storability-related quantitative trait loci (QTLs) discovered in previous studies, suggesting that these loci may provide important regions for isolating the responsible genes. In total, 448 annotated genes were predicted within the identified regions, of which 274 and 82 had nonsynonymous and frameshift mutations, respectively. We detected extensive metabolic activities and cellular processes during seed storability and confirmed the effects of the seed storability-related candidate loci using four BILs from DXWR and R974. These results may facilitate the cloning of DXWR seed storability-related genes, thereby elucidating rice seed storability and its improvement potential.

## 1. Introduction

Rice (*Oryza sativa* L.) plays a vital role in food production and security globally [1]. However, rice seeds can lose their viability and vigour during storage very easily, which has caused serious yield losses. Seed aging and the decrease of vigour during storage have been the main problems of rice production [2,3]. In recent years, the problem of seed storage has become increasingly prominent in China owing to the continuous harvest of rice, especially in the humid and hot south area [4,5]. Currently, seed aging is delayed mainly by improving the storage conditions, which is costly and energy-consuming [6]. Breeding rice cultivars having strong seed storability is one of the most cost-effective and efficient ways to alleviate this problem.

Seed storability is a complex quantitative trait and is affected by both external and internal factors [2,7]. Nowadays, external factors, such as chemical agents and the manipulation of temperature and humidity, are mainly used to prolong storage time; however, these processes are costly and rice seeds may be contaminated easily [6]. Therefore, it is of great significance to study how internal factors can be used to improve the seed storability of rice. The internal factors include the genetic background, fatty acid content, starch characteristics, antioxidants, etc. A number of physiological and biochemical studies have revealed that many changes can affect seed vigour, including membrane system, DNA, and protein damage; decreased protein synthesis ability; and the accumulation of excessive reactive oxygen species [8,9]. Meanwhile, the loss of three kinds of lipoxygenases (Lox1, Lox2, and Lox3) could delay the aging and deterioration of rice seeds, resulting in improved tolerance to storage [6,10].

During the past decades, more than 70 quantitative trait loci (QTLs) related with seed storability have been identified in different rice accessions [11]. However, the traditional QTL analysis relies on the development of mapping populations, which is time-consuming and easily affected by genetic background noise [12]. Meanwhile, most of the QTL analyses on seed storability have been performed on populations derived from modern rice varieties, and most of the detected QTLs were isolated in large intervals, making subsequent fine-scale mapping difficult. Furthermore, the genetic resources of seed storability in modern rice varieties are limited, and some valuable genes of rice seed were lost during artificial selection and the long-term evolution process of rice [13]. Therefore, utilizing genetic resources from other rice species, especially in wild rice relatives, is significant for rice breeding strategies for seed storability improvement.

The advancement of next generation sequencing (NGS) technology provides an opportunity for exploring the genetic diversity among various rice accessions and its utilization in breeding programs. For example, Liang et al. applied NGS technology to identify a novel *pi21* haplotype conferring basal resistance to rice blast disease [14]. Kitony et al. used NGS technology for population development and days to heading (DTH) QTLs mapping [15]. Reyes et al. utilized NGS technology for marker-assisted backcross breeding to introgress and stack the *Gn1a* and *WFP* alleles to improve rice cultivars [16]. On the other hand, bulked segregant analysis via whole genome sequencing (BSA-seq) is a fast and powerful tool to accurately identify QTLs associated with complex traits [17]. BSA-seq has been widely used in various plants, such as rice [18], cucumbers [19], and cotton [20]. However, there is limited research on the identification of elite QTLs from wild rice by BSA-seq. The Dongxiang wild rice, found in Dongxiang county, Jiangxi province, China, is the northernmost (28°14’N) wild rice species globally. DXWR possesses many elite genetic resources related with tolerance against various abiotic stresses, such as cold, drought, salt, and hot temperature [21]. Additionally, DXWR has strong seed storability and harbours many valuable genetic resources [22]. However, the molecular mechanisms of the seed storability of DXWR remain poorly understood. Therefore, the main aim of the present study was to identify and analyse seed storability-related loci in DXWR using BSA-seq, which will be helpful in the fine-scale mapping and cloning of novel genetic resources improving the seed storability of rice.

## 2. Materials and Methods

### 2.1. Plant materials

The parents DXWR and F6, and the derived backcross inbred lines (BILs) populations (BC_2_F_7_; 252 lines) were used in the BSA-seq. Another BIL population (BC_4_F_6_; 84 lines) was derived from DXWR and R974. Using molecular marker background analysis, four BILs (BIL27, BIL49, BIL64, and BIL203) from the DXWR/R974 population were selected to analyse the effects of the candidate loci. In the two populations, the receptor parents, F6 and R974, are modern rice varieties and the common donor parent DXWR is an accession of *Oryza rufipogon*. The two populations were cultivated in the paddy fields of Jiangxi Normal University, Nanchang city, China in the summer of 2017. The seeds of the populations and parents were harvested, dried, and stored in a refrigerator at −20 °C.

### 2.2. Seed storability evaluation and extreme bulk sample construction

For seed storability evaluation, the seeds were initially subjected to a 20-day artificial aging treatment in the dark at 42 °C, in an environment with 88% humidity [11]. The treated seeds were then placed in a petri dish with a filter paper, were kept moist using sterile water, and then incubated in a constant temperature incubator at 32 °C. The germination standard was based on the length of the germinated radicle or embryo reaching half the length of the seed [23]. After 5 days, the germination rates were counted. The germination rates (%) = number of germinated seeds/total number of seeds × 100%. For BSA-seq, 20 lines with high germination rates and 20 lines with low germination rates were selected to construct bulked-tolerant (TB-bulk) and bulked-intolerant (IB-bulk) BIL samples, respectively [24]. The experiment was repeated three times to ensure the correct phenotype. Statistical analysis was performed using Student’s *t*-test.

### 2.3. Sequence libraries construction and BSA-seq

Sequence libraries construction and BSA-seq were performed by Biomarker Technologies Corporation (Beijing, China). The procedure was carried out according to the protocol provided by Illumina (San Diego, CA, USA). Briefly, the qualified DNA was randomly fragmented into 350 bp length by ultrasonic processor. Then, the DNA fragments were end repaired, and a A-tail was added to the 3’ end, followed by adapter ligation and purified. Subsequently, the sequencing libraries were constructed by PCR amplification and sequenced by Illumna Novaseq 6000. The raw image data obtained by sequencing were transformed by base calling software (Illunima Casava v.1.8.2) and stored in FASTQ format. Then, the raw reads were filtered to obtain clean reads for further bioinformatics analysis. The clean reads from four libraries were separately mapped to the Nipponbare MSU v7 reference genome (http://rice.uga.edu/) using BWA software [25]. BSA-seq was conducted as previously described [26]. All SNPs and InDels variations were identified by Unified Genotype function of the Genome Analysis Toolkit (GATK) software [27], and the SnpEff software [28] was used to annotate the SNPs and InDels.

For the results of BWA alignment, use Picard (http://sourceforge.net/projects/picard/) to de-duplication and shield the influence of PCR-duplication; use GATK for local re-alignment, base quality re-calibration and variant calling. The detection of SNP/InDel was mainly performed by the GATK software toolkit [27]. Raw SNPs were filtered with GATK according to the following criteria: filter out if there were 2 SNPs within 5 bp, SNP within 5 bp near InDel was filtered; two InDel distances less than 10 bp are filtered out. Before performing association analysis, we filtered again in order to obtain high quality SNP/InDel sites [29]. SNP-index and InDel-index of each SNP and InDel position were calculated to identify the candidate loci associated with seed storability. To minimize false-positive sites, we used the positions of markers in the genome to fit the ΔSNP-index and ΔInDel-index values marked on the same chromosome using the DISTANCE method [30]. Additionally, a Euclidean distance (ED) measurement algorithm was also applied as previously described [31].

BLAST software [32] was used to annotate the genes in the identified candidate loci for seed storage tolerance with NT, NR [33], Swiss-Prot [34], GO [35], KEGG [36] and COG [37] databases. All annotated genes containing nonsynonymous and frameshift mutations in the identified candidate loci associated with seed storage tolerance were selected for further analysis.

### 2.4. Molecular marker background analysis

Total genomic DNA was extracted from fresh leaves by the Plant Genomic DNA Extraction Kit (Sangon Biotech, Shanghai, China). PCR system and amplification procedure were performed as previously described [38]. PCR products were separated on 3% agarose gels. SSR marker sequences were found in the Gramene website (www.gramene.org), and InDel marker sequences were designed by Primer3 Input (version 0.4.0) (https://bioinfo.ut.ee/primer3-0.4.0/). All of the molecular markers were synthesized by Sangon Biotech Company Limited (Shanghai, China) (Table S1).

## 3. Results

### 3.1. Identification of BILs with extreme phenotypes for seed storability

After the artificial aging treatment, the germination rate of DXWR was approximately 42.50%, while that of F6 was 18.33%. The germination rate differences between DXWR and F6 were significant (Figure 1a,b). Meanwhile, for each BIL of the population derived from DXWR and F6, the germination rates had obvious differences, ranging from 0% to 57.50%, with an average value of 19.20% (Figure 1c). The distribution of the germination rates of the BILs had a skewed normal distribution. There were 118 BILs (46.83%) had higher germination rates than that of the receptor parent F6, suggesting that an extensive introgression of genes from DXWR to F6 occurred in the population. Then, 20 BILs with high germination rates were grouped as TB-bulk sample and 20 BILs with low germination rates were grouped as IB-bulk sample for further BSA-seq analysis.

**Figure 1.**
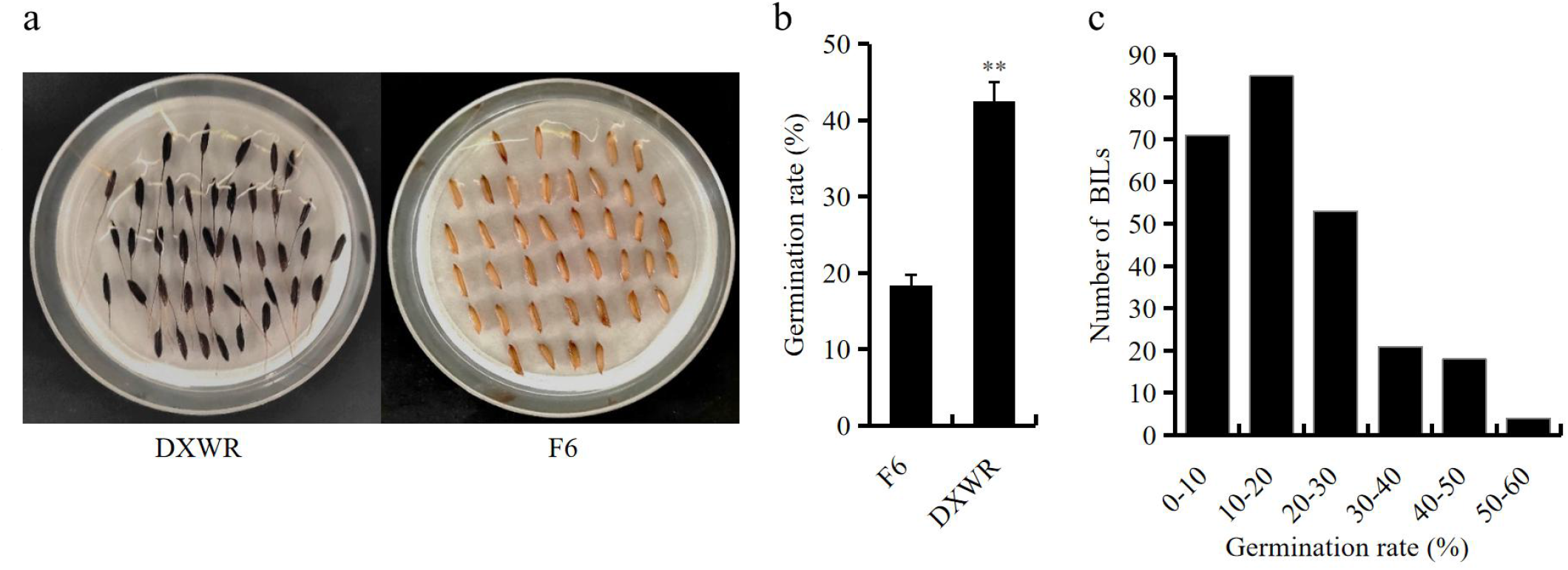
Germination rates of DXWR, F6 and their BILs after aging treatment. (**a**) the germination rate of DXWR was significantly higher than that of F6 after aging treatment. (**b**) average germination rates of DXWR and F6 after aging treatment. Asterisks mark significant differences according to Student’s *t*-test, **P value < 0.01. (**c**) distribution of germination rates in BILs population from crosses DXWR and F6 after aging treatment.

### 3.2. BSA-seq, data analysis, and quality assessment

Using whole genome sequencing for the libraries of DXWR, F6, TB-bulk, and IB-bulk, we generated 33,922,707, 33,982,607, 39,408,702, and 33,710,561 raw reads, respectively (Table 1). After filtering, we obtained a total of 42.19 Gb of clean data, including 10.09 Gb for IB-bulk and 11.79 Gb for TB-bulk. The total amounts of clean data for the donor parent DXWR and the receptor parent F6 were 10.16 Gb and 10.14 Gb, respectively. In total, approximately 96.32% of the clean reads could be aligned to the reference genome. In this study, the average GC content of the four samples was 44.56%, thereby indicating that the GC distribution was normal. Meanwhile, the average depth of the genome coverage of the four samples was 21.50 ×, while the average genomic coverage was 94.36% at 1× coverage and 89.28% at 5 × coverage. In addition, the *Q*-value (*Q*30) was used as the cut-off for quality control. The *Q*30 values of all samples were > 90%, ranging from 91.54% (TB-bulk) to 94.70% (F6), with an average of 93.56% (Table 1). These results indicated that the data were of good quality and could be used for subsequent analysis.

**Table 1.** The quality of sequencing data and analysis of sequencing depth and coverage.

|  | Raw reads | Clean base (bp) | Mapping rate (%) | *Q*30 (%) | GC (%) | Average depth (×) | Coverage at least 1× (%) | Coverage at least 5× (%) |
| --- | --- | --- | --- | --- | --- | --- | --- | --- |
| DXWR | 33,922,707 | 10,157,432,212 | 97.54 | 93.95 | 44.63 | 20 | 97.36 | 93.27 |
| F6 | 33,982,607 | 10,144,910,192 | 96.60 | 94.70 | 45.32 | 21 | 91.06 | 86.93 |
| IB-bulk | 33,710,561 | 10,092,480,184 | 95.82 | 94.03 | 44.58 | 21 | 94.59 | 88.27 |
| TB-bulk | 39,408,702 | 11,791,891,500 | 95.31 | 91.54 | 43.71 | 24 | 94.44 | 88.63 |

### 3.3. Identification of candidate genome loci for seed storability

After filtering, a total of 330,715 high-quality SNPs and 74,076 high-quality InDels were identified among the different samples. In general, the greater the ΔSNP-index and ΔInDel-index, the more likely the contribution of SNPs and InDels to the trait of interest or their association with the candidate genes that control the trait. In this study, the ΔSNP-index and ΔInDel-index graphs were plotted and computed against the genome positions (Figure 2). The SNP analysis indicated that the regions containing four genomic loci on chromosome 4 and three genomic loci on chromosome 9 could be the candidate loci associated with seed storability. The seven loci covered a total length of 3.65 Mb and contained a total of 548 genes (Table S2). On the other hand, the InDel analysis indicated that the regions containing five genomic loci on chromosome 4 and one genomic locus on chromosome 9 could contribute to the trait of seed storability. These six loci covered a total length of 3.49 Mb and contained a total of 516 genes (Table S2). Several candidate loci at the intersection of the SNP and InDel analysis were identified at the 18,550,000–20,870,000 bp on chromosome 4 (named as locus-4) and 7,860,000–9,780,000 on chromosome 9 (named as locus-9) and are considered as candidate loci for seed storability of DXWR (Table 2). Within the identified regions, a total of 448 annotated genes were identified (Table 2). Among them, 274 genes had nonsynonymous mutations and 82 genes had frameshift mutations (Table S3).

**Table 2.** Candidate regions of seed storability of DXWR.

| Chromosome ID | Start | End | Size (MB) | Gene number |
| --- | --- | --- | --- | --- |
| Chr4 | 18,550,000 | 18,550,000 | 0.00 | 1 |
| Chr4 | 18,600,000 | 19,670,000 | 1.07 | 162 |
| Chr4 | 20,620,000 | 20,630,000 | 0.01 | 3 |
| Chr4 | 20,650,000 | 20,680,000 | 0.03 | 3 |
| Chr4 | 20,770,000 | 20,820,000 | 0.05 | 8 |
| Chr4 | 20,860,000 | 20,870,000 | 0.01 | 1 |
| Chr9 | 7,860,000 | 9,780,000 | 1.92 | 270 |
| Total | - | - | 3.09 | 448 |

*3.4. Bioinformatics analysis of annotated genes in the identified regions*

**Figure 2.**
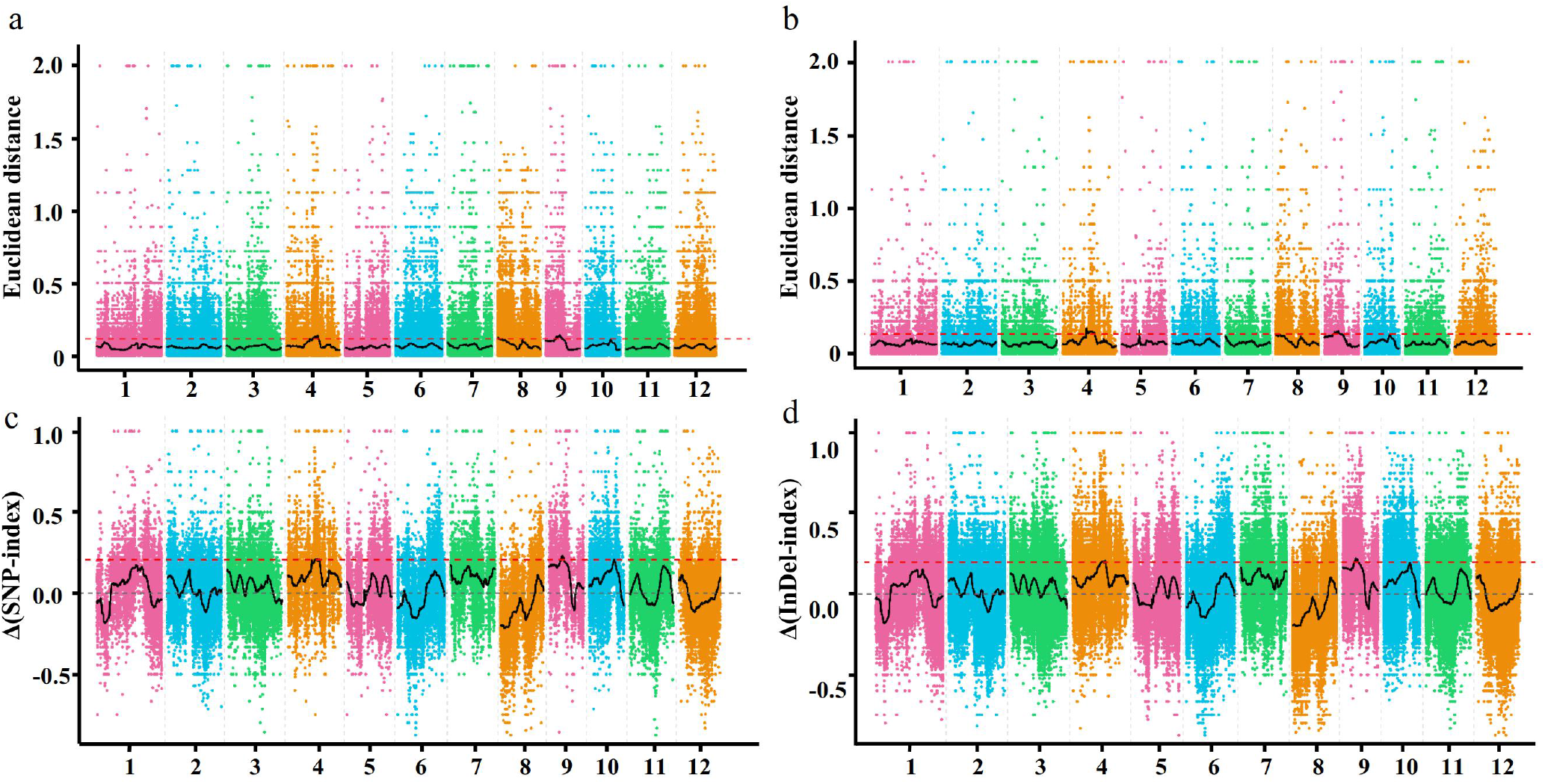
The identifcation of candidate genome regions for seed storability by Euclidean Distance (ED) and delta-index methods. (**a**) SNP-ED association algorithm. (**b**) InDel-ED association algorithm. (**c**) Δ(SNP-index) method. (**d**) Δ(InDel-index) method. The abscissa is the chromosome number, the colored dot represents the calculated ED value or Δ(SNP-index)/Δ(InDel-index) value of the SNP/InDel site, and the black line is the fitted ED value or Δ(SNP-index)/Δ(InDel-index) value. The red dashed horizontal lines represent the signifcance association threshold.

### 3.4. Bioinformatics analysis of annotated genes in the identified regions

To analyse the annotated genes in the identified regions, we carried out bioinformatics analysis using multiple databases. As shown in Table 3, according to the NR database, the candidate regions contained 431 annotated genes, with 265 genes having nonsynonymous mutations and 78 genes having frameshift mutations. However, based on the NT database, we found 448 annotated genes in the candidate loci. Among them, 274 genes had nonsynonymous mutations and 82 genes had frameshift mutations. Meanwhile, we analysed the annotated genes in the Swiss-Prot database and found that 194 genes could be annotated. Among them, 116 genes had nonsynonymous mutations and 28 genes had frameshift mutations. In addition to Swiss-Prot database, we also analysed genes using the TrEMBL database. In total, we found 443 genes in the candidate loci that could be annotated on the TrEMBL database. Among them, 267 genes had nonsynonymous mutations and 78 genes had frameshift mutations. Furthermore, we identified 337 annotated genes in the candidate loci based on the GO database. Among them, 196 genes had nonsynonymous mutations and 53 genes had frameshift mutations.

**Table 3.** Information of annotated genes in the identified regions using multiple databases.

| Annotated databases | Gene number | Nonsynonymous- mutation gene number | Frameshift- mutation gene number |
| --- | --- | --- | --- |
| NR | 431 | 265 | 78 |
| NT | 448 | 274 | 82 |
| TrEMBL | 433 | 267 | 78 |
| SwissProt | 194 | 116 | 28 |
| GO | 337 | 196 | 53 |
| KEGG | 64 | 32 | 4 |
| GOG | 97 | 61 | 15 |
| Total | 448 | 274 | 82 |

Further, the annotated genes were sorted into GO term categories. The results illustrated that the annotated genes could be clustered into 26, 65, and 74 GO terms in the cellular component (Table S4), molecular function (Table S5), and biological process categories (Table S6), respectively. Within the cellular component category, the cell, organelle, and cell parts were the most abundant. Within the molecular function category, catalytic activity and binding were the most highly represented groups. Metabolic processes and cellular processes were the most abundant in the biological processes category (Figure S1, Table S7). Meanwhile, the annotated genes were also found to be enriched in other stress-related terms, such as biological regulation, stimulus response, nucleic acid binding transcription factor, and transporter activity (Figure S1).

Next, the functional hierarchy of the annotated genes in the KEGG orthology system was classified into five categories, including cellular processes, environmental information processing, genetic information processing, metabolism, and organismal systems. We found annotated 64 genes in the candidate loci based on the KEGG database. Among them, 32 genes had nonsynonymous mutations and four genes had frameshift mutations (Table S3). The statistical results showed that most of the annotated genes were associated with metabolism, followed by genetic information processing and cellular processes (Figure S2). Moreover, a KEGG reference pathway analysis showed that the top five enriched pathways were lysine degradation, citrate cycle (TCA cycle), tryptophan metabolism, C5-branched dibasic acid metabolism, and peroxisome (Table S8).

### 3.5. Confirmation of the candidate loci for seed storability

Using molecular marker background analysis, four BILs (BIL27, BIL49, BIL64, and BIL203) from the DXWR and R974 population were selected to confirm the effect of the candidate loci. In BIL27 and BIL64, segments of chromosome 9 including the candidate locus-9 were substituted with those of DXWR in the background of R974 and no segments including the candidate locus-4 were substituted (Figure 3). In addition, the segments of chromosome 4 including the candidate locus-4 were substituted in BIL49 and BIL203 (Figure 3). Meanwhile, BIL49 and BIL203 had no segments including the candidate locus-9 of DXWR (Figure 3). After an artificial aging treatment, the germination rate of R974 was found to be 23.33%, which was significantly lower than that of DXWR. Meanwhile, the germination rates of BIL27, BIL49, BIL64, and BIL203 were 57.50%, 54.17%, 61.67%, and 50.83%, respectively (Figure 4). Therefore, the germination rates of the four BILs carrying the candidate loci of the seed storability of DXWR were significantly higher than those of R974, implying that the candidate loci could contain the causative genes associated with seed storability in DXWR.

**Figure 3.**
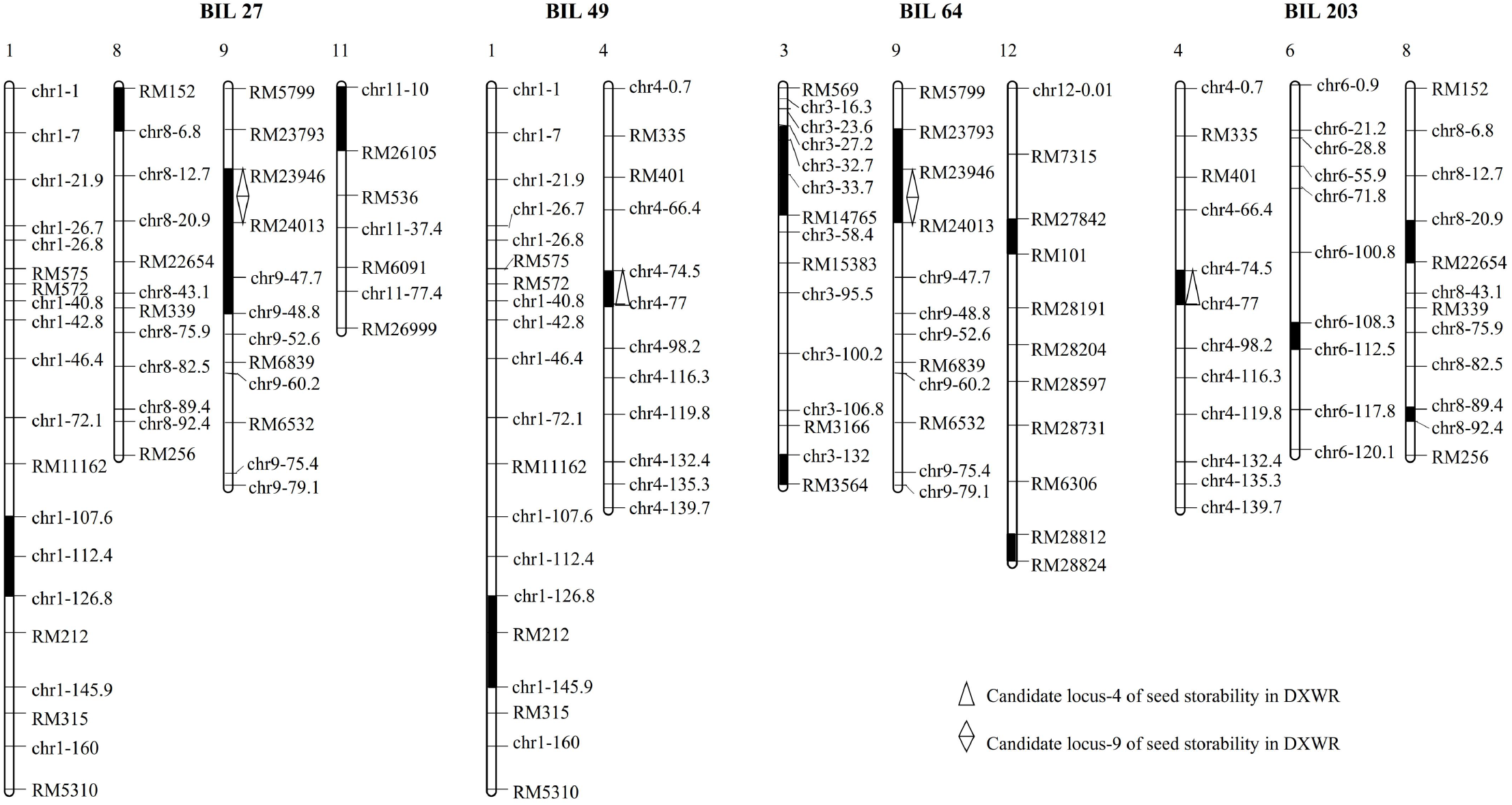
Graphical representation of four BILs (BIL27, BIL49, BIL64, and BIL203) from the DXWR and R974 population. White squares represent homozygous for R974 allele, black squares represent homozygous for DXWR allele.

**Figure 4.**
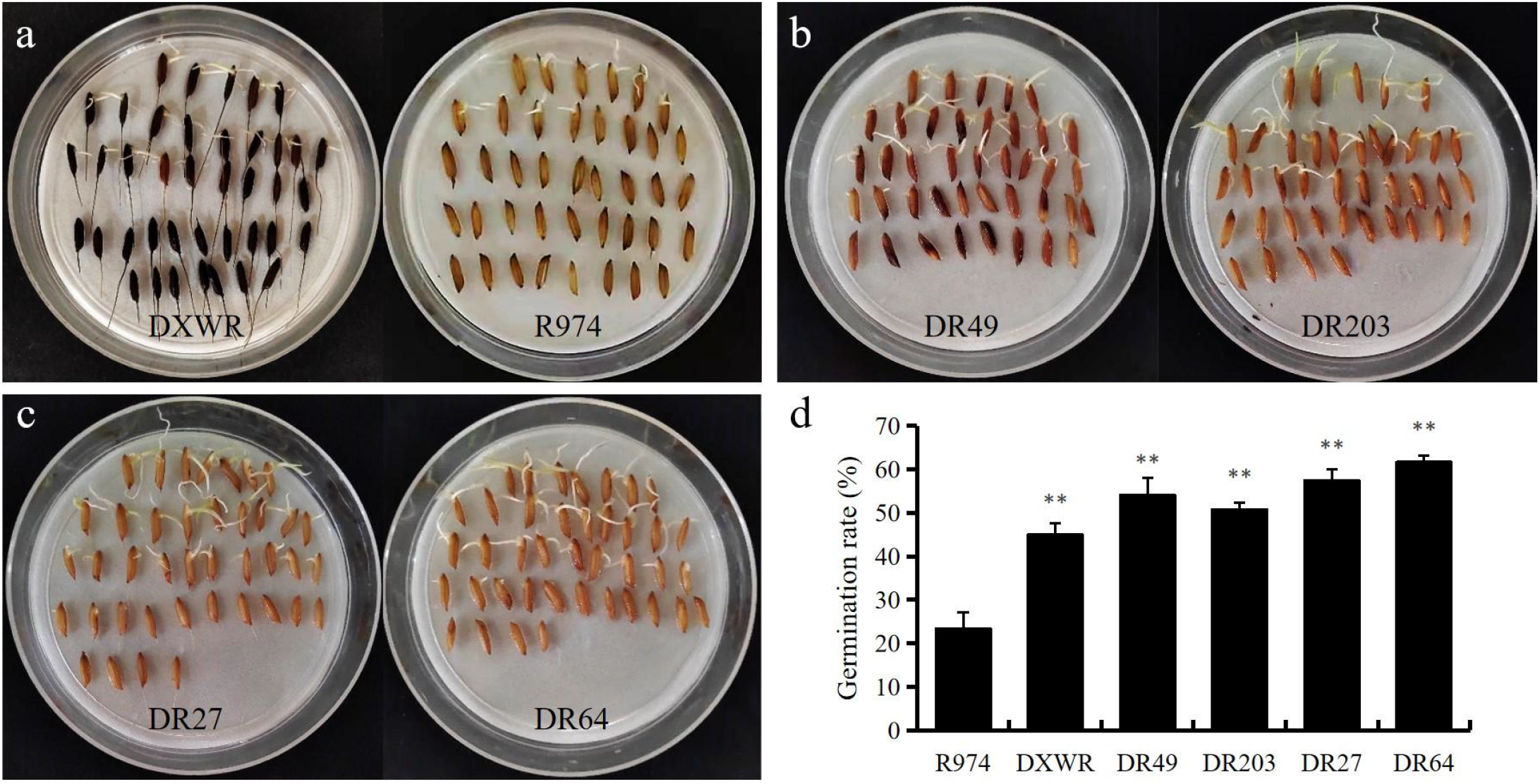
Germination rates of the parents and the four BILs after aging treatment. (a) Comparison of germination rates between DXWR and R974 after aging treatment. (**b**) phenotype chart of DR49 and DR203. (**c**) phenotype chart of DR27 and DR64. (**d**) average germination rates of parents and four BILs after aging treatment. Asterisks mark significant differences according to Student’s *t*-test, **P value < 0.01.

## 4. Discussion

In agronomy, the seed storability of rice is an important trait owing to its correlation with seed quality and germination [39]. Thus, understanding the mechanisms of seed storability and breeding varieties with high storability is critical for rice production. Seed storability is a complex quantitative trait that is controlled by large numbers of QTLs [40]. We recently developed IncRNA-derived-SSR markers and the SSR markers from drought stress-responsive miRNAs [49, 50]. Although some progress has been made in traditional QTL mapping, the underlying molecular mechanisms of seed storability in rice remain poorly defined. Identifying more seed storability-related QTLs from different rice accessions is becoming quite valuable. DXWR has strong seed storability and is an ideal rice material to identify seed storability-related QTLs and illustrate the molecular mechanisms [22]. In this study, we identified the candidate genomic regions on chromosomes 4 and 9 for the seed storability of DXWR. Our results have greatly narrowed the interval for identifying the seed storability-related genes of DXWR and have provided valuable information for the fine-scale mapping and cloning of the seed storability-related genes.

The degree of resistance to aging varies among different rice varieties, which is restricted by nongenetic and genetic factors [41]. Till now, a number of studies have identified many QTLs associated with seed storability in different rice accessions; some of these QTLs were also located on chromosomes 4 and 9. For example, using a set of 85 BILs derived from a cross between Sasanishiki and Habataki, Hang et al. identified a total of 13 seed storability-related QTLs on chromosomes 1, 2, 3, 4, 5, 7, 11, and 12. Among these QTLs, *qSSh-4* was mapped between two markers (R288 and C891) on chromosome 4; the existence of *qSSh-4* was further confirmed by Habataki chromosome segment substitution lines in a Sasanishiki genetic background [42]. Using two different segregating populations, Lin et al. identified seven QTLs for seed storability on chromosomes 1, 2, 5, 6, and 9. The QTL *qSSn-9* was simultaneously detected at almost the same region in the interval between RM444 and RM105 on chromosome 9 in the two populations, implying that *qSSn-9* was a stable QTL for seed storability [39]. Meanwhile, Li et al. identified 10 QTLs for seed germination percentage using a double haploid population derived from the cross between CJ06 and TN1 during either natural storage or artificial aging. Among the identified QTLs, *qGP-9* on chromosome 9 was delimited to an interval of 92.8 kb between two STS markers (P6 and P8) [43]. Furthermore, in some studies, the seed storability-related QTLs were also identified on chromosomes 4 and 9 simultaneously. For example, using 98 BILs derived from a cross between Nipponbare and Kasalath, Miura et al. detected three QTLs for seed longevity, including *qLG-2, qLG-4*, and *qLG-9* on chromosomes 2, 4, and 9, respectively, after applying an aging treatment. *qLG-4* was located between markers C1100 and C1016 on chromosome 4, and *qLG-9* was located between markers C103 and C1751 on chromosome 9 [44]. Subsequently, Sasaki et al. fine mapped *qLG-9* into a 30-kb interval of the Nipponbare genome sequence [45]. Using a BIL population derived from a cross between Koshihikari and Kasalath, Li et al. detected six QTLs for seed storability on chromosomes 2, 3, 4, 6, 9, and 11. Among them, *qSS-4* was located on chromosome 4 between markers C513 and C1100; *qSS-9* was located on chromosome 9 between markers R10783S and S752 [46]. By performing a QTL comparative analysis, the candidate locus-4 and locus-9 identified in our study appeared to partially overlap with *qSSh-4, qSS-9*, and *qSSn-9*. These results imply that these loci could be specific and reliable seed storability-related loci.

A total of 448 annotated genes were identified within our identified loci, of which 274 genes had nonsynonymous mutations and 82 genes had frameshift mutations. The bioinformatics analysis of these annotated genes revealed extensive metabolic activities and cellular processes during seed storability. An in-depth analysis of the biological process annotation of GO revealed that *LOC_Os09g13610* was involved in the biological process of jasmonic acid (JA) mediated signalling pathway (GO:0009867). Previous studies had revealed that the hormone JA is a critical regulator of tolerance to abiotic stresses [47,48]. In addition, we found that two genes (*LOC_Os04g32020* and *LOC_Os09g14670*) were involved in the tricarboxylic acid cycle process (GO:0006099), one gene (*LOC_Os04g32070*) was involved in the oxidation-reduction process (GO:0055114), and one gene (*LOC_Os09g15670*) was involved in the abiotic stress responses (GO:0009409, GO:0009414). Interestingly, out of the five annotated genes identified in the aforementioned GO database, only *LOC_Os09g13610* was also annotated by the KEGG database. Furthermore, the sequence alignment results showed that the locus of *LOC_Os09g13610* had many nonsynonymous or frameshift mutations, which differentiated the parents as well as the two bulked pools. These annotated genes could be involved in the regulation of seed storability and need further investigation.

Using genetic background analysis, we selected four BILs from the DXWR and R974 population to analyse the effect of locus-4 and locus-9. The results showed that the germination rates of the BILs harbouring the identified loci of seed storability were significantly higher than those of R974 after implementing the artificial aging treatments. Therefore, we concluded that the candidate loci identified in our study could contain the genes related to the seed storability of DXWR; additionally, the tightly linked markers of the loci will be valuable in future breeding programs to develop new cultivars with improved rice seed storability.

## 5. Conclusions

In this study, we applied the BSA-seq method and identified the seed storability-related loci of DXWR. By performing a comparative analysis of QTLs, we found that the candidate loci overlapped partially with the seed storability-related QTLs found in previous studies. Meanwhile, 448 annotated genes were identified within the candidate loci, and large numbers of nonsynonymous and frameshift mutations existed in these genes. Additionally, the candidate loci were confirmed by four BILs from the DXWR and R974 population. These results will provide genetic information on the cloning and molecular marker-assisted selection of the seed storability-related genes of DXWR, and thus will be helpful in advancing the breeding strategies and breeding resources required to increase the seed storability of rice.

## Supplementary Materials

Figure S1: GO enrichment analysis of annotated genes in the candidate regions. The ordinate is each GO category, the abscissa is the number of genes. Figure S2: KEGG enrichment analysis of annotated genes in the candidate regions. The abscissa is the number of genes annotated to the pathway, and the ordinate is the name of KEGG metabolic pathway. Figure S3: Venn diagram of SNP and InDel loci detection for parental DXWR and F6 and TB-bulk and IB-bulk samples. Table S1: The molecular markers used in graphic genotyping of the BILs population of DXWR and R974. Table S2: SNP- and InDel-index association results. Table S3: Nonsynonymous and frameshift mutations in the annotated genes of the identified regions. Table S4: GO analysis in the cellular component category of annotated genes in the identified regions. Table S5: GO analysis in the molecular function category of annotated genes in the identified regions. Table S6: GO analysis in the biological process category of annotated genes in the identified regions. Table S7: GO enrichment analysis of annotated genes in the identified regions. Table S8: KEGG enrichment analysis of annotated genes in the candidate regions.

## Author Contributions

M.Z., B.H. and G.D. performed the experiments. M.Z. and W.Y. analyzed the data. Y.H.C. and Y.C. helped in the sample collection. Y.F. helped to edit the manuscript. M.Z. B.H., Y.F. and F.Z. drafted the manuscript. F.Z. and J.X. contributed to the experimental design and edition of the manuscript. All authors have read and agreed to the published version of the manuscript.

## Institutional Review Board Statement

Not applicable.

## Informed Consent Statement

Not applicable.

